# Combining agronomic biofortification with a Zn-efficient wheat line synergistically increases grain Zn concentrations with varied effects on Cd in wheat grains

**DOI:** 10.64898/2026.10.18.759907

**Authors:** Jill Bachelder, Julie Tolu, Lenny H.E. Winkel, Navreet K. Bhullar, Ingrid Stoffel-Studer, Chloe Manzanares, Steven Andrew Yates, Matthias Wiggenhauser, Emmanuel Frossard

**Affiliations:** Group of Plant Nutrition, Institute of Agricultural Sciences, Department of Environment Systems; ETH Zurich, Lindau, Switzerland; Group of Inorganic Environmental Geochemistry, Institute of Biogeochemistry and Pollutant Dynamics, Department of Environment Systems, ETH Zurich, Zurich, Switzerland; Department of Water Resources and Drinking Water, Eawag, Dübendorf, Switzerland; Plant Molecular Biology, Department of Biology, ETH Zurich, Zurich, Switzerland; Molecular Plant Breeding, Institute of Agricultural Sciences, ETH Zurich, Zurich, Switzerland

**Keywords:** wheat, biofortification, micronutrients, agriculture, zinc, cadmium

## Abstract

**Background and aims:** Wheat grains are often too low in zinc (Zn) compared to human dietary needs, and are a major dietary source of highly toxic cadmium (Cd). Biofortification strategies can increase Zn in edible grain, but must not increase grain Cd.

**Methods:** We investigated the individual and combined effects of genetic and agronomic biofortification on Zn and Cd in wheat grains. Non-transgenic Bobwhite (BW-NT) and an *OsNAS2-*expressing line (BW-NAS) were grown in a pot experiment with isotope-labeled mineral Zn (^70^Zn) and green manure (^70^Zn, ^106^Cd). At flowering and maturity, Zn and Cd were quantified in wheat biomass. Wheat contents of Zn and Cd derived from inputs versus soil were determined using stable isotope analyses.

**Results:** Individually, the *OsNAS2-*expressing wheat and input application increased grain Zn. Combined *OsNAS2* expression and input application led to grain concentrations of 76±4 mg kg^−1^ with mineral Zn application and 61±7 mg kg^−1^ with green manure. This was 154-217% higher than the 24±1 mg kg^−1^ observed in the BW-NT control and significantly higher than the additive effects of each strategy individually. The *OsNAS2-*expressing line increased grain Cd to 0.214±0.002 mg kg^−1^ under green manure, compared to 0.12±0.01 mg kg^−1^ in the BW-NT control. No other treatment increased grain Cd.

**Conclusions:** Our results encourage development of integrated genetic and agronomic biofortification strategies to increase grain Zn. They also underscore the need for careful monitoring of grain Cd concentrations when organic amendments are paired with genetic biofortification strategies.

## Introduction

It is estimated that 46% of the global population does not consume sufficient dietary zinc (Zn) (Passarelli et al. 2024). A major cause of insufficient dietary Zn intake is low contents and poor availability of Zn in cereal-derived food products (Graham et al. 2007; Cakmak and Kutman 2018). To combat Zn deficiency, biofortification strategies can be used to increase Zn contents in edible plant products (Frossard et al. 2000; Bouis and Welch 2010). Such strategies include genetic biofortification, e.g. plant breeding and bioengineering techniques; agronomic biofortification, e.g. application of inputs to increase soil Zn; or a combination thereof (Frossard et al. 2000). Yet, biofortification strategies (especially genetically engineered) are rarely evaluated for their effects on the highly toxic element cadmium (Cd) (Singh et al. 2017; Cakmak and Kutman 2018; Nayak et al. 2023, 2025), despite the fact that wheat can accumulate high concentrations of Cd and is a major source of chronic exposure (EFSA 2012; Zhao et al. 2023). Furthermore, Zn and Cd have similar biological uptake pathways in wheat, with increased grain Zn in cereal crops shown to be associated with increased Cd (Detterbeck et al. 2016). Thus, it is essential to investigate whether there exist genetic and agronomic biofortification strategies that can increase Zn without increasing Cd.

The genetic biofortification technique of constitutively expressing the *Nicotianamine synthase (NAS)* gene has been demonstrated to increase grain Zn concentrations in various cereal crops (Wirth et al. 2009; Masuda et al. 2009; Johnson et al. 2011; Kawakami and Bhullar 2018, 2020; Lee et al. 2023), including wheat (Singh et al. 2017), but the effects on grain Cd concentration are largely unknown. The *NAS* genes encode enzymes responsible for production of nicotianamine, a metal chelating molecule that plays an important role in Zn transport and storage within plants, e.g. in roots and grains (Johnson et al. 2011; Haydon et al. 2012; Clemens et al. 2013). Nicotianamine is the main chelator of Zn in the phloem (Nishiyama et al. 2012; Olsen and Palmgren 2014) and is involved in xylem loading from the roots (Cornu et al. 2015). Constitutive of *NAS* also leads to increased production of the phytosiderophore 2’deoxymugineic acid (DMA) (Nozoye 2018), as nicotianamine is a precursor of DMA (Shojima et al. 1990). Root release of DMA occurs under low iron (Fe) or Zn availability to solubilize previously unavailable micronutrients (Cakmak et al. 1996; Oburger et al. 2014), and also plays a role in Zn transport and storage (Kobayashi et al. 2019; Clemens 2019). Chelators such as DMA have been shown to selectively promote the uptake of Zn micronutrients over toxic elements such as Cd (Hill et al. 2002; Banakar et al. 2017). This is due to their higher binding affinity for micronutrients like Zn and Fe over Cd (Shenker et al. 2001; Hill et al. 2002; Puschenreiter et al. 2017). However, it has not yet been investigated whether constitutive expression of *NAS* genes in wheat could lead to a selective increase in Zn uptake without enhancing Cd uptake.

Genetic biofortification strategies can be complemented by agronomic strategies such as input application to increase soil available Zn (i.e., forms that can be taken up by plants). Application of mineral Zn fertilizers, e.g. ZnSO_4_, increases Zn uptake in wheat without increasing Cd (Hart et al. 2002). This occurs via the addition of available Zn that competes with Cd for uptake (Hart et al. 2002; Cakmak 2008; Zou et al. 2012; Rehman et al. 2021). The few previous studies investigating combined genetic and agronomic biofortification have focused on application of water-soluble mineral Zn fertilizers, with promising increases of 20-25% in grain Zn concentration compared to genetic biofortification alone (Nayak et al. 2023, 2025). In contrast, there are very few studies of combined genetic biofortification with organic amendment application, despite the fact that organic amendments can also increase soil available Zn (Watson et al. 2006; Grüter et al. 2017; Bachelder et al. 2025). Organic amendments can stimulate Zn uptake in wheat not only via addition of available Zn species to soil, but also by promoting solubilization of native soil pools, e.g. following the degradation of organic matter (Zou et al. 2012; Aghili et al. 2014; Grüter et al. 2017; Künzli et al. 2024). Due to the high complexity of these processes and variability in the chemical composition of different organic amendments, organic amendments show varied abilities to increase grain Zn concentrations (Soltani et al. 2014; Helfenstein et al. 2016; Grüter et al. 2019; Bachelder et al. 2026). Via the same processes, organic amendments may also increase Cd concentrations in wheat grains (Schweizer et al. 2018). Due to the wide application of organic amendments and their overall positive effects on soil fertility (Dobermann et al. 2022), it is important to investigate how genetic biofortification can be combined not only with water-soluble mineral Zn but also with organic amendment application.

We investigated the individual and combined effects of a single *OsNAS2-*expressing line and input application (water-soluble mineral Zn, green manure) on Zn and Cd concentrations in wheat grains. We hypothesized that the constitutive expression of *OsNAS2* would increase Zn concentrations in grains without increasing Cd, due to higher affinity of N- and O-containing organic ligands (such as nicotianamine and DMA) for binding Zn than Cd (Suzuki et al. 2007) and evidence of preferential wheat uptake of phytosiderophore-Zn complexes over Cd complexes (Shenker et al. 2001). We also expected combined *NAS* constitutive expression and input application would lead to even higher Zn concentrations in wheat grains but not Cd, due to the high affinity of DMA for binding Zn, which would allow increased uptake with higher availability in soil. To test these hypotheses, we performed a pot experiment in which a non-transgenic and an *OsNAS2*-expressing Bobwhite wheat line were grown in an arable, non-contaminated soil. To distinguish between soil native pools and input-derived elements, we applied water-soluble mineral Zn labeled with ^70^Zn and green manure labeled with ^70^Zn and ^106^Cd. The results of this study provide insight into whether biofortification techniques may be combined to produce wheat grains with optimum Zn concentrations, without increasing Cd.

## Materials & methods

### Soil collection and characterization

Soil collection and characterization has been described previously (Schweizer et al. 2024; Bachelder et al. 2026). In summary, soil was collected at a depth of 0-20 cm from an agricultural field in Jura, Switzerland, followed by air-drying. A subsample was sieved to 2 mm and analyzed by Sol-Conseil for pH (water, 1:2.5), cation exchange capacity, and texture (Agroscope 2020a, b, c). For further analyses and pot experiments, soil was passed through a 6 mm plastic sieve. Water-holding capacity was determined at ambient pressure (Agroscope 2020d). Total carbon and nitrogen were determined with an elemental analyzer (Vario PYRO cube, Elementar GmbH, Germany). Diethylenetriaminepentaacetic acid (DTPA)-extractable Zn and Cd were determined via extraction following the protocol of Lindsay and Norvell (1978) and analysis with inductively coupled plasma mass spectrometry (ICP-MS; Agilent 7500), with instrumental details provided in <u>Bachelder et al. (2026)</u>. DTPA-Zn was 0.8 mg Zn (kg soil)^−1^, falling in the range <1 mg (kg soil)^−1^ that may indicate low Zn availability in soil (Brennan et al. 1993). Concentrations of Zn and Cd were determined by microwave acid digestion with HF following the procedure of Imseng et al. (2019) and quantification via ICP-MS. Soil properties were published previously in Bachelder et al. (2026) and Schweizer et al. (2024), and are provided in Table S1. All values throughout this manuscript, including Table S1, are provided in dry weight.

### Production of isotope-labeled inputs and Bobwhite wheat seeds

Metallic ^70^Zn (97.4 atom% relative abundance) and ^106^Cd (99.5 atom% relative abundance) were obtained from CortecNet SA (Les Ulis, France) and digested in 8 M HNO_3_ (double-distilled from Supelco EMSURE®). To prepare isotope-labeled ZnSO_4_, digestate was diluted to 400 mg ^70^Zn (L)^−1^ and mixed 1:10 with a ZnSO_4_ solution of 400 Zn mg (L)^−1^. This resulted in a solution with ^70^Zn abundance of 10% and ^66^Zn abundance of 25%, as confirmed by isotope analyses (see “Plant Analyses”). Isotope-labeled green manure (sunflower) was produced in hydroponics containing full-strength Hoagland solution (Hoagland and Arnon 1950). The solution was spiked with ^70^Zn and ^106^Cd prepared from 50:50 mixtures with non-labeled Zn(NO_3_)_2_ and Cd(NO_3_)_2_ solutions, respectively (Table S2). The produced sunflower contained ^70^Zn abundance of 26%, ^66^Zn abundance of 21%, ^106^Cd abundance of 49%, and ^111^Cd abundance of 7% as confirmed by isotope analyses (see “Plant Analyses”). Total elemental analysis of C and N in plants was performed as for soil. Mineral N was determined by extraction with 0.01 M CaCl_2_ (Sigma-Aldrich, 99.99% trace metals basis; solid-liquid 1:20) followed by NH_4_^+^ quantification with Berhelot’s reaction (Searle 1984) and NO_3_^−^quantification by ion chromatography (Metrohm, 930 Compact IC Flex with chemical suppression). The chemical composition of the isotope-labeled green manure is provided in Table S1.

Seeds of non-transgenic wheat cultivar Bobwhite S26 (hereafter BW-NT) and an *OsNAS2-*expressing Bobwhite line (hereafter BW-NAS) were obtained from the authors of Singh et al. (2017). A detailed description of the protocol for production of the transgenic wheat line has been reported previously (Singh et al. 2017). Singh et al. (2017) demonstrated that all *OsNAS2* single-copy insertion lines showed increased Fe and Zn accumulation, indicating that the phenotype is robust across independent events. As we performed a full-factorial pot experiment (two wheat lines x three agronomic treatments x two destructive harvests), it was not experimentally feasible to include more than one *OsNAS2-*expressing wheat line. Therefore, only line 65 (T4 generation) was chosen, as this was shown to have the highest expression of *OsNAS2* out of all transformed lines and significantly increased Fe and Zn concentrations in the grains compared to the non-transgenic line. Because only one line was used, we highlight that the current study serves as a novel investigation exploring the possibilities of combining genetic and agronomic biofortification strategies, and does not aim to provide full proof for commercial implementation of *NAS* overexpression combined with input application. A new generation of seeds (i.e., T5) was produced for this study by growing the seeds of BW-NT and BW-NAS in five pots of ∼2 kg potting soil. During seed production, the new generation of BW-NT and BW-NAS were fertilized with eight applications of nutrient solution (Table S3) and grains were harvested at maturity, i.e., five months after sowing.

### Pot experiment

To compare with our previous pot experiment (Bachelder et al. 2026), 200 μg Zn (kg soil)^−1^ and 2 μg Cd (kg soil)^−1^ in the form of non-labeled Zn(NO_3_)_2_ and Cd(NO_3_)_2_, respectively, were added to each 1 kg of soil. This was considered as part of the “native” soil pool (i.e., not from inputs) due to the twelve-week aging process after application and homogeneous application across all pots regardless of treatment. Further explanation is provided in Note S1. After 12 weeks of incubation, a basal nutrient solution was applied to the soil in addition to three input treatments: no Zn input (hereafter, “control”), ZnSO_4_ labeled with ^70^Zn (hereafter, “mineral Zn”), and green manure labeled with ^70^Zn and ^106^Cd. The basal nutrient solution was adapted in the green manure treatment to maintain a constant application rate of mineral N in all treatments (Table S4). Seeds of BW-NT and BW-NAS were sterilized for 15 minutes in 30 % H_2_O_2_ solution, rinsed with ultrapure water, and germinated in triple-washed 2 mm sand. Seedlings were sown one-per-pot three days after input and nutrient solution application, to create six treatments in total: BW-NT grown under control, mineral Zn, and green manure and BW-NAS grown under control, mineral Zn, and green manure.

Wheat was grown for four weeks in a greenhouse set to 14 h photoperiod (65% relative humidity, 23 °C) and 10 h darkness (60% relative humidity, 20 °C). Pesticides were sprayed four weeks after sowing due to presence of powdery mildew on leaves. The pesticides contained 0.5 mL L^−1^ “Input” (Bayer, active agents: spiroxamine, Prothioconazole) and 0.3 mL L^−1^ “Etalfix Pro” (Syngenta, active agent: polyether modified trisiloxane). The total concentrations of Zn and Cd in applied pesticide mix were quantified using ICP-MS and were found to be 114 μg (L)^−1^ Zn and 1.5 μg (L)^−1^ Cd. After spraying, wheat was moved to a growth chamber for the remainder of the experiment. Growth chamber settings included 13 h photoperiod (illuminance of 25 klx, 55 % relative humidity, 24° C) and 11 h darkness (50 % relative humidity, 18 °C). During plant growth, soil was watered by weight to maintain 30-70% water-holding capacity. Pots were randomized once per week. Eight experimental (pot) replicates were included per treatment. Half of the replicates were harvested at flowering (87 days after sowing) and remaining pots were harvested at maturity (129 days after sowing). At harvests, aerial biomass (which we define as all aerial organs, i.e., stem, leaves, husks, and grains) was cut 2 cm above the soil surface and rinsed with ultrapure water. At maturity, grains were separated from aerial biomass. Root washing was performed in three successive baths of 4 °C 6 mM NaCl, for a total duration of 10 minutes (Böhm 1979). After harvesting, all biomass was frozen and freeze-dried.

### Plant analyses

Confirmation of *OsNAS2* expression was performed via reverse transcription-polymerase chain reaction (RT-PCR) following the workflow reported previously (Singh et al. 2017). Leaves of BW-NT and BW-NAS plants were collected at flowering and flash-frozen in liquid N_2_. Single biological replicates (BW-NT with green manure, BW-NAS with mineral Zn) were processed in triplicates. RNA was extracted using the Nucleospin RNA Plus kit (Macherey-Nagel). RNA samples were treated with DNAse (ThermoFisher, DNA-free™ DNA Removal Kit) followed by cDNA synthesis (ThermoFisher, RevertAid First Strand cDNA Synthesis kit). Primers were designed to target the *OsNAS2* gene (GCAAGTGCTGCAAGATGG forward, CCGGATCAGACGGATAGC reverse). The gene *Ta.22845* was used as a reference gene to validate the gene expression (AAGAAGCAAGAGCTCGGATTT forward, TGCACCGTTGAAATCATCA reverse) (Singh et al. 2017). Both genes were amplified using GoTaqG2 DNA Polymerase and the following PCR settings: 95°C for 2 min.; 35 cycles of 95°C for 30 sec., 60°C for 30 sec., 72°C for 20 sec.; 72°C for 5 min. Gel electrophoresis was performed to confirm the expression of *OsNAS2* in BW-NAS and not BW-NT.

For phenotypic characterization of BW-NT versus BW-NAS wheat lines, “days to flowering” was recorded as the stage when at least three spikes per plant had produced flowers. The 1000 grain weight was determined by weighing three replicates of 50 seeds and extrapolating to 1000. Plant height, ears per plant, grain biomass, and total biomass were determined at maturity.

For chemical characterization, plant samples were ground with a tungsten-carbide ball mill (Qiagen TissueLyzer). Total elemental analysis of C, N, and S was performed as for soil. Plant samples (200 mg) were digested in 4 mL of 8 M HNO_3_ using a single-reaction chamber microwave system (turboWave, MWS), as reported previously (Bachelder et al. 2026). Metal quantification was performed using triple-quadrupole ICP-MS/MS (Agilent 8900), with all instrument parameters reported previously (Bachelder et al. 2026). As a quality control, samples were digested and analyzed in parallel with a WEPAL certified reference material (Table S5). Elemental concentrations were calculated per mass of plant samples. Total elemental contents were calculated by multiplying elemental concentrations by plant biomass.

In treatments receiving labeled inputs, isotope ratios (^70^Zn/^66^Zn and ^106^Cd/^111^Cd) were determined using a modified method of McBeath et al. (2013), as reported in our previous study (Bachelder et al. 2026). Briefly, digested plant samples were purified to remove interfering matrix elements (Dürr-Auster et al. 2019; Yan et al. 2021; Künzli et al. 2024). Samples were transferred to acid-cleaned PTFE beakers, evaporated to dryness, redissolved in 0.3 M HCl, and loaded onto PTFE mini-columns (Sun et al. 2021) containing anion exchange resin (AG 1-X8, 100–200 mesh, chloride form, Bio-Rad laboratories) conditioned with 1 M HCl. Columns were washed with 1 M HCl to remove matrix elements, followed by Zn and Cd elution with 0.3 M HNO_3_. The Agilent 7500 ICP-MS was used to measure stable isotopes of Zn (^64^Zn, ^66^Zn, ^67^Zn, ^68^Zn, ^70^Zn), Cd (^106^Cd, ^108^Cd, ^110^Cd, ^111^Cd, ^112^Cd, ^113^Cd, ^114^Cd, ^116^Cd), and all isobaric interfering elements (^60^Ni, ^72^Ge, ^105^Pd, ^115^In, and ^115^Sn). Every four samples, a natural-abundance bracket solution was analyzed that contained 40 µg Zn (L)^−1^ and 2 µg Cd (L)^−1^. Repeated analysis of bracket standards (n = 20) showed relative standard deviations of 1% for ^66^Zn, 2% for ^70^Zn, 2% for ^111^Cd, and 6% for ^106^Cd. Blanks contained <5% Zn and Cd compared to samples. Mass bias correction of isotopes was calculated following Equation (1) for Zn and Equation (2) for Cd, with mass difference *M* (calculated as *y-66* for Zn, *y-111* for Cd) and mass bias *k* calculated following Tolu et al. (2014).

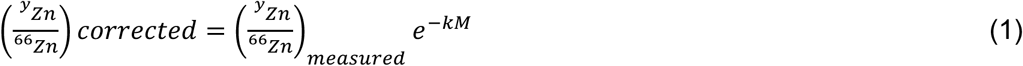

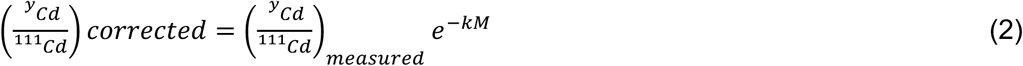

The fractions of Zn and Cd in plant biomass derived from inputs were determined following Equations (3) and (4), respectively. The abundances of ^66^Zn, ^70^Zn, ^111^Cd, and ^106^Cd were determined in the labeled mineral Zn and green manure inputs (A_66,Input_, A_70,Input_, A_111,Input_, A_106,Input_). Soil abundances of ^66^Zn, ^70^Zn, ^111^Cd, and ^106^Cd were set to natural abundance values of IUPAC (A_66,Soil_, A_70,Soil_, A_111,Soil_, A_106,Soil_) (Berglund and Wieser 2011). Isotope ratios in plant biomass were considered as the “sink”.

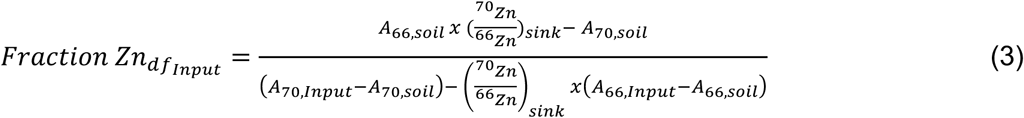

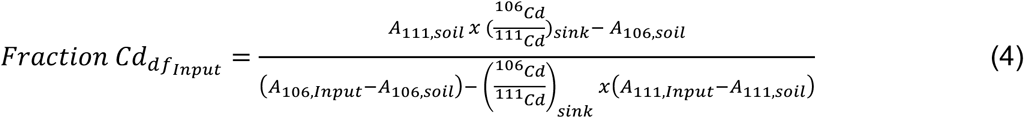

The total mass of input-derived Zn and Cd at maturity was calculated by multiplying the fractions determined by Equations (3) and (4) with the total contents of Zn and Cd in wheat biomass. These values were used to determine the use efficiencies (i.e., recoveries) of Zn and Cd added as inputs using Equations (5) and (6), respectively. Zn_sink_ indicates the Zn mass in plant biomass (i.e., the entire wheat plant). Zn_input_ indicates the mass of Zn added to a pot with the input application (i.e., with mineral Zn or green manure).

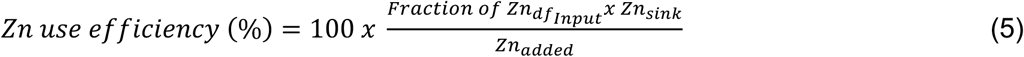

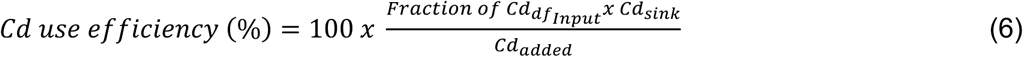

### Soil analyses

Soil was sampled destructively during harvest at flowering. Analysis of pH was performed in ultrapure water (1:2.5). Samples were extracted with ultrapure water (1:10), filtered to 0.45 μm (BGB, Nylon 66), and analyzed for dissolved organic carbon (Shimadzu, TOC-L). Dissolved metals in water extracts were quantified using ICP-MS/MS, with NIST 1643f certified reference material as quality control (Table S6).

### Calculations and statistics

To evaluate internal translocation between plant organs, transfer factors were calculated as concentration in the sink organ(s) divided by concentration in the source organ(s). Statistical tests were performed using R studio (version 2022.02.3+492). Generalized least squares and emmeans post hoc tests were used to investigate significant differences between input treatments and wheat lines (confidence level = 0.95). For response variables deemed heteroscedastic by the Levene test, significant differences and interactions were determined using generalized least squares modeling with the “varIdent” constant variance function. Significant interactions of treatments (wheat line, inputs, or line*inputs) were determined using two-way ANOVA calculations using the generalized least squares model output. As Cd source tracing was only performed for one input treatment (green manure), statistical differences were determined using one-way analysis of variance and Tukey post hoc tests. A paired t-test was performed using Microsoft Excel (version 16.90) to compare the total uptake of Zn and Cd at harvest 1 with harvest 2, as these data could be considered to be dependent (i.e., uptake at harvest 2 is linked to uptake at harvest 1). An independent t-test was used to compare the increase in Zn grain concentrations and total uptake between (1) the combined *OsNAS2* constitutive expression and input treatments and (2) the additive effects of each treatment individually. Independent t-tests were calculated based on propagated errors of sums and differences between values in, e.g., BW-NAS/mineral Zn and BW-NT/control, following equations provided by Ross and Willson (2017). For all comparisons, statistically significant differences were accepted when *p*-values were less than 0.05.

## Results

Expression of *OsNAS2* in BW-NAS was confirmed by RT-PCR, which also showed no expression in BW-NT (Figure S1). The RT-PCR passed our quality control, by showing expression of the *Ta.22845* reference gene in both lines. The BW-NAS line showed slight phenotypic differences compared to BW-NT. Compared to the BW-NT control, the BW-NAS control plants were 14% taller (Figure 1B), which is expected as reported in the original work by <u>Singh et al. (2017</u>). Only in the green manure treatment, BW-NAS produced significantly less grain biomass than the BW-NT line (Figure 1E). Otherwise, we found no statistically significant differences in phenotypic metrics (Figure 1).

**Figure 1.**
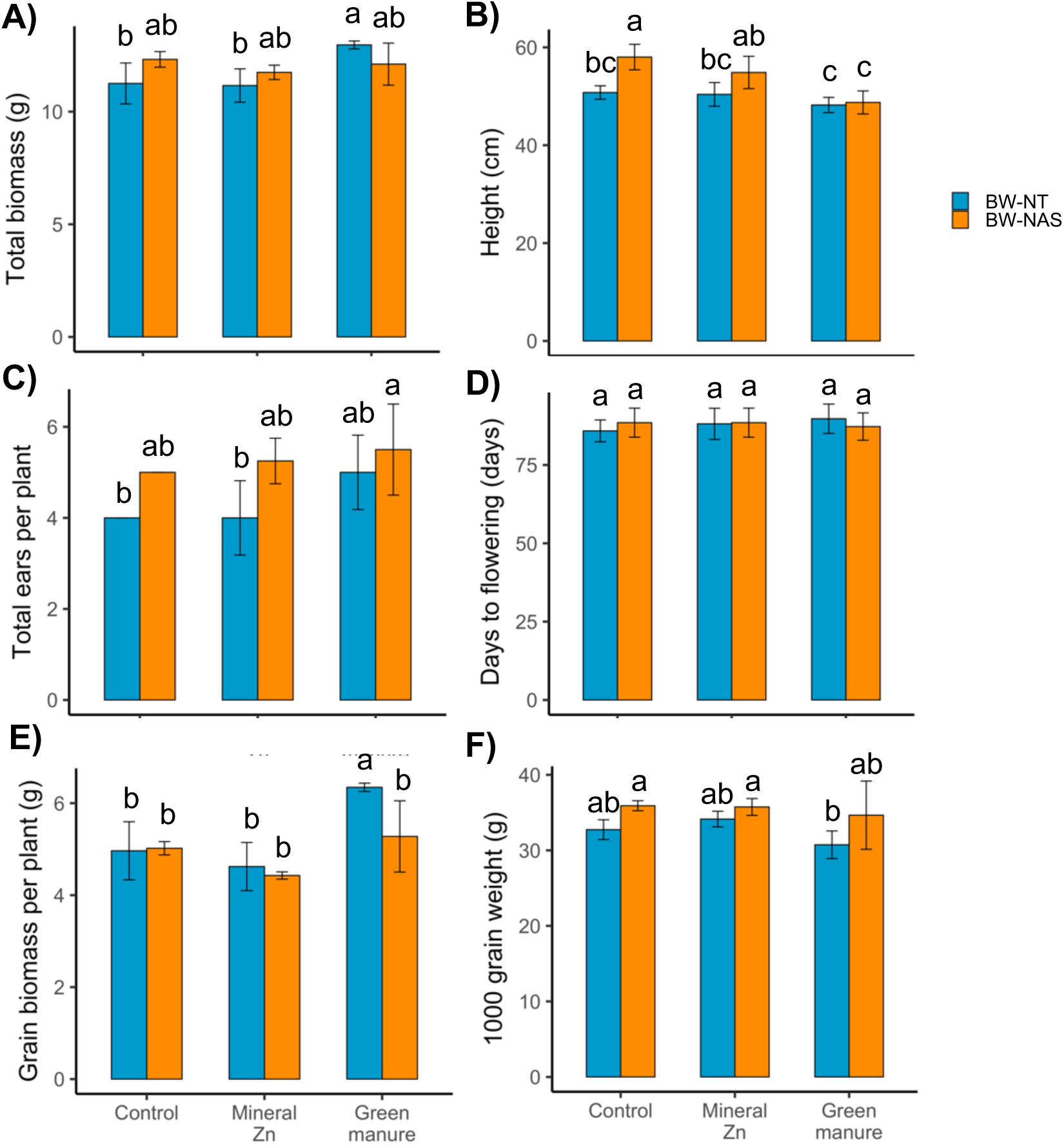
Phenotypic metrics for comparing *OsNAS2-*expressing Bobwhite (BW-NAS) and non-transgenic (BW-NT) wheat lines. Analyzed parameters included A) total wheat biomass per plant at full maturity, B) plant height at full maturity, C) number of ears per plant at maturity, D) days to flowering, E) grain biomass per plant, and F) 1000 grain weight. Plants of BW-NT and BW-NAS were grown under three input treatments (control, mineral Zn, and green manure application). Error bars indicate one standard deviation about the mean, of n=4 biological (pot) replicates. Lowercase letters indicate statistical differences calculated using generalized least squares modelling followed by emmeans posthoc tests. Significant *p*-values for the two-way ANOVA analyses based on generalized least squares output were as follows: A) inputs *p*=0.03, line*inputs *p*=0.003; B) line *p*<0.001, inputs *p*<0.001, line*inputs *p*=0.02; C) line *p*<0.001, inputs *p*=0.0498; D) ; E) line *p*=0.03, inputs *p*<0.001, line*inputs *p*=0.04; and F) line *p*=0.001.

To investigate the combined effect of constitutive *OsNAS2* expression and input application on wheat biofortification, we analyzed total grain concentrations of Zn and Cd. We found grain Zn increased in BW-NAS compared to BW-NT by a factor of 2, to 51±1 mg Zn (kg grain)^−1^ from 24±1 mg Zn (kg grain)^−1^ in the control treatment (Figure 2A). In both lines, mineral Zn application led to the highest grain Zn concentrations, though this increase was only statistically significant in the BW-NAS line. Green manure application also increased grain Zn in both lines, though this was less than for mineral Zn and again only statistically significant in the BW-NAS line. Our data also provided the opportunity to compare the individual and combined effects of our wheat lines and input treatments. Mineral Zn application led to an increase of 8 mg Zn (kg grain)^−1^ in BW-NT and 32 mg Zn (kg grain)^−1^ in BW-NAS compared to the respective control treatments. Yet, the difference between BW-NAS grown with mineral Zn and BW-NT grown under the control was 52 mg Zn (kg grain)^−1^, which is higher than the sum of each treatment individually, i.e., 40 mg Zn (kg grain)^−1^. Similarly, growing an *OsNAS2*-expressing wheat line with green manure led to higher grain Zn than the expected additive effect of each treatment individually, i.e., 38 mg Zn (kg grain)^−1^ observed compared to 17 mg Zn (kg grain)^−1^ additive. A t-test based on propagated errors confirmed the combined increases to be significantly higher than the additive increases for mineral Zn (*p*<0.001) and green manure (*p*=0.04).

**Figure 2.**
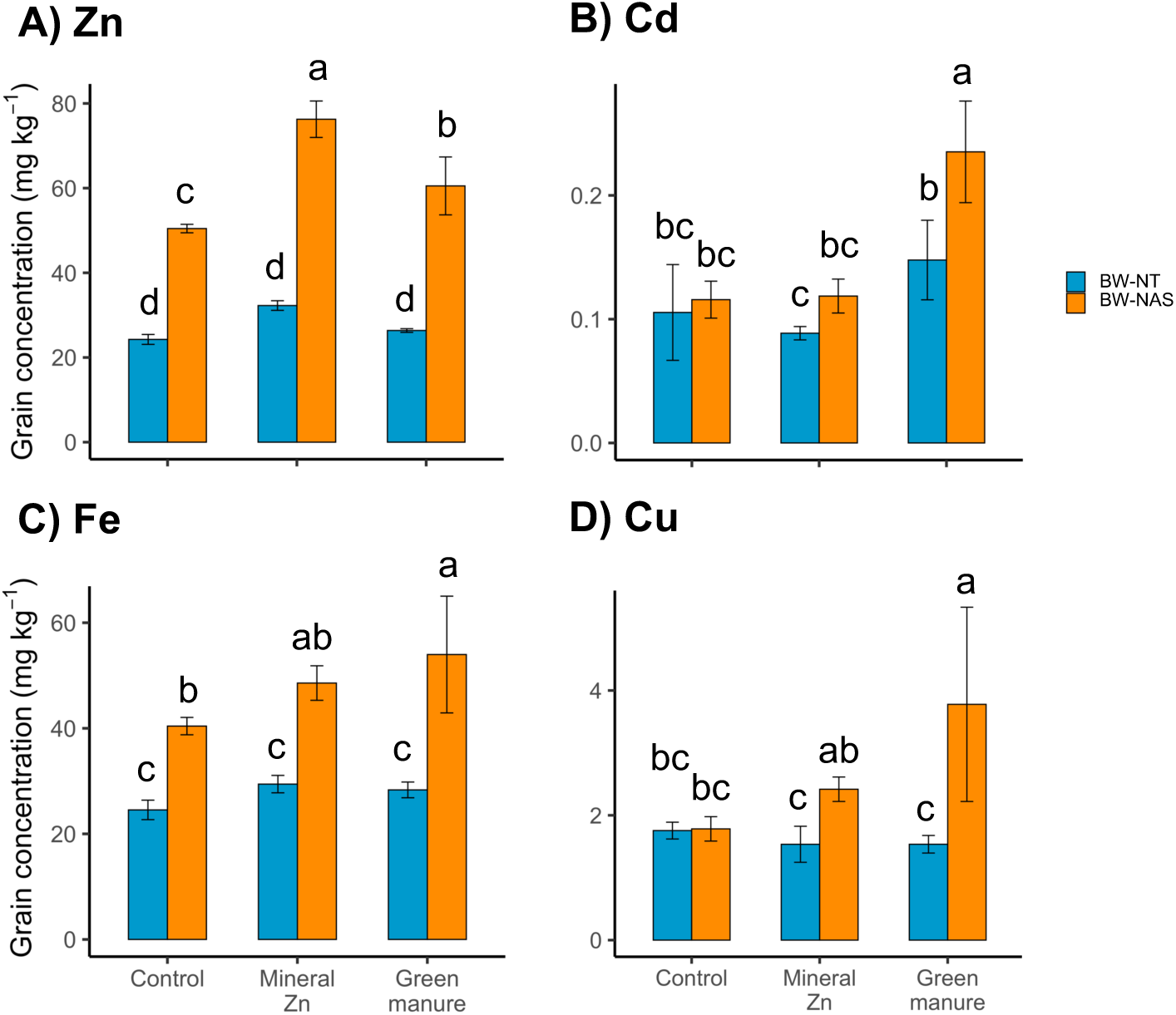
Grain concentrations of A) Zn, B) Cd, C) Fe, and D) Cu measured in non-transgenic (BW-NT) and *OsNAS2*-expressing (BW-NAS) wheat lines grown under three input treatments (control, mineral Zn, and sunflower green manure application). Error bars indicate one standard deviation about the mean, of n=4 biological (pot) replicates. Lowercase letters indicate statistical differences calculated using generalized least squares modelling followed by emmeans posthoc tests. Significant *p*-values for the two-way ANOVA analyses based on generalized least squares output were as follows: A) line *p*<0.001, inputs *p*<0.001, line:inputs *p*<0.001; B) line *p*<0.001, inputs *p*<0.001, line:inputs *p*=0.02; C) line *p*<0.001, inputs *p*=0.002; D) line *p*<0.001, inputs *p*=0.02, line*inputs *p*=0.006.

In contrast to Zn, no increase in grain Cd was observed in BW-NAS in the control treatment compared to BW-NT (Figure 2B). In the BW-NT line, no inputs led to a significant difference in grain Cd compared to the control treatment. Green manure increased grain Cd by a factor of 2 in the BW-NAS line, to 0.214±0.002 mg Cd (kg grain)^−1^ from 0.12±0.01 mg Cd (kg grain)^−1^ in the BW-NT control. BW-NAS contained higher grain Fe in all input treatments compared to BW-NT (by a factor of 1.6-1.9; Figure 2C). We also observed significantly higher grain Cu with mineral Zn (factor 1.6) and green manure application (factor 2.5) compared to the respective BW-NT lines (Figure 2D). Concentrations of other elements did not vary between wheat lines (Figure S2).

To better understand the factors controlling grain concentrations, we first investigated how wheat line and input treatments affected uptake of Zn and Cd in the entire plant (i.e., roots plus aerial biomass) at flowering and maturity. Under all treatments, BW-NAS took up approximately 100 μg Zn (kg soil)^−1^ between flowering and maturity, whereas no significant Zn uptake was observed in BW-NT between flowering and maturity (Table S7). Similar to grain concentrations, BW-NAS showed two-times higher Zn uptake, i.e. 391±32 μg Zn (kg soil)^−1^ in compared to 185±5 μg Zn (kg soil)^−1^ in BW-NT at maturity (Figure 2A). Mineral Zn increased wheat Zn uptake at maturity by a factor of 1.5 in BW-NAS compared to the BW-NAS control (Figure 3A). Green manure application did not alter Zn uptake in either wheat line compared to the respective controls. The expected additive effect of the *OsNAS2-*expressing line and mineral Zn, i.e., 250 μg Zn (kg soil)^−1^, was much lower than the observed increase of 409 μg Zn (kg soil)^−1^ in BW-NAS with mineral Zn compared to the BW-NT control. This followed the same pattern as observed for grain concentrations. However, the additive effect of the *OsNAS2-*expressing line and green manure application, i.e., 256 μg Zn (kg soil)^−1^, was not far from the observed combined increase of 217 μg Zn (kg soil)^−1^. A t-test confirmed the combined increases to be significantly higher than the additive increases for mineral Zn (*p*=0.001) but not green manure (*p*=0.07). No treatment led to a statistically significant change in Cd uptake (Figure 3D). BW-NAS showed increased Fe uptake in all input treatments by a factor of 1.5 and increased Cu with green manure by a factor of 1.2, compared to the BW-NT controls (Figure S3-D,E).

**Figure 3.**
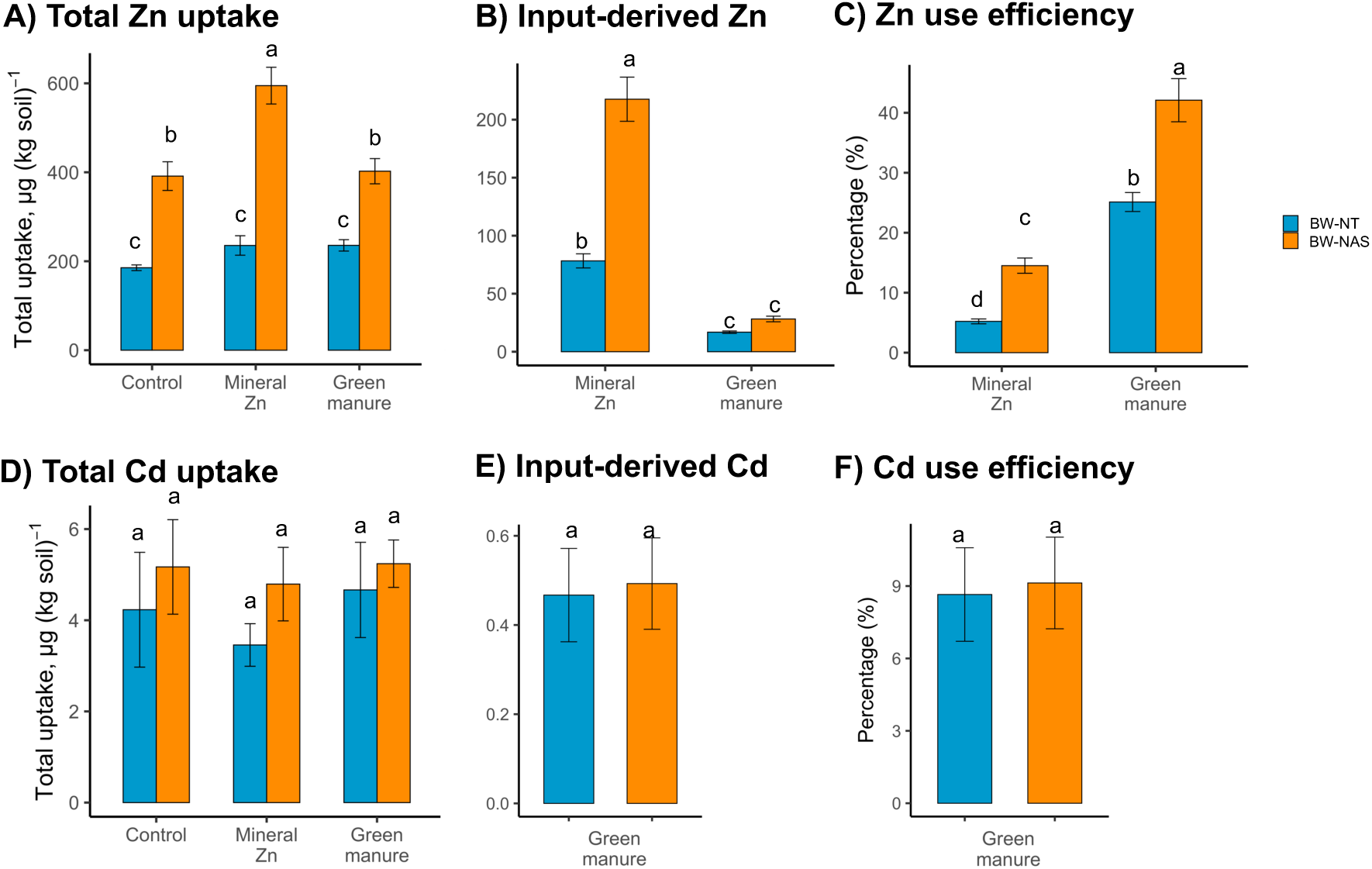
Total and input-derived uptake of Zn and Cd in non-transgenic (BW-NT) and *OsNAS2*-expressing (BW-NAS) wheat lines grown under three input treatments (control, mineral Zn, and sunflower green manure application). Total uptake was determined at maturity by summing elemental contents in roots and all aerial biomass (stem, leaves, and grains). Input-derived values were determined by multiplying total uptake by the fraction of input-derived Zn and Cd calculated via isotope labeling. Use efficiency was calculated as the percentage of Zn or Cd applied to the soil that was quantified as input-derived at maturity. Error bars indicate one standard deviation about the mean, of n=4 biological (pot) replicates. Lowercase letters indicate statistical differences calculated using generalized least squares modelling followed by emmeans posthoc tests. Significant *p-*values for the two-way ANOVA analyses based on generalized least squares output were as follows: A) line *p*<0.001, inputs *p*<0.001, line*inputs *p*<0.001; B) line *p*<0.001, inputs *p*<0.001, line*inputs *p*<0.001; C) line *p*<0.001, inputs *p*<0.001, line*inputs *p*<0.001; and D) line *p*=0.01.

We then investigated whether soil chemical properties relevant for metal availability (pH, DOC, water-soluble Zn, Cd, Fe, Cu; Figure S4) could explain differences in plant uptake of Zn and Cd. An increase in soil water-soluble Cu was observed for BW-NAS with green manure, compared to the BW-NT control (Figure S4-C). Otherwise, no difference in soil chemical parameters was observed between any treatment and the BW-NT control. Thus, increases in Zn and Fe total uptake in BW-NAS wheat compared to BW-NT wheat were not explained by soil chemical parameters.

Stable isotope tracing allowed us to distinguish the sources of Zn and Cd that were taken up by wheat (i.e., between input- and soil-derived pools). From the mineral Zn treatment, the BW-NT control took up 78±6 μg Zn (kg soil)^−1^ input-derived Zn, which increased by a factor of 2.8 to 217±19 μg Zn (kg soil)^−1^ in BW-NAS (Figure 3B). With green manure, BW-NT took up 17±1 μg Zn (kg soil)^−1^ input-derived Zn, which increased to 28±2 μg Zn (kg soil)^−1^ in BW-NAS. In contrast, there was no statistically significant difference in input-derived Cd uptake between BW-NT and BW-NAS (Figure 3E). No input treatment led to a difference in soil-derived Zn or Cd uptake compared to the respective BW-NT/BW-NAS controls (Figure S5).

We calculated use efficiency (i.e., recovery) by comparing the application rates of Zn and Cd in our treatments (Table S4) and the input-derived contents in plant biomass (Figure 3B,E). With mineral Zn, Zn use efficiency was 5.2±0.4 % in BW-NT and 15±1 % in BW-NAS (Figure 3C). With green manure, Zn use efficiency was 25±2 % in BW-NT and 42±4 % in BW-NAS. Thus, Zn use efficiency was consistently higher with green manure than mineral Zn and higher in BW-NAS than BW-NT lines. In contrast to Zn, Cd use efficiency was the same in BW-NT and BW-NAS lines, ranging from 7-12 % (Figure 3F). Comparison of these results with our previous experiment is provided in Table S8 and Note S2.

To investigate internal translocation of Zn and Cd, we calculated transfer factors based on tissue concentrations at maturity. Transfer factors from roots to aerial biomass ranged from 1-5 for Zn and 0.1 to 0.4 for Cd (Figure 4A,C). We observed increased transfer of Zn from roots to aerial biomass in BW-NAS lines compared to BW-NT, by a factor of 2-3 across all input treatments (Figure 4A). With green manure, we observed increased transfer of Zn from non-grain aerial biomass to grains in the BW-NAS line compared to the BW-NT control (Figure 4B). Otherwise, no treatment significantly altered transfer to grains. In contrast to Zn, no treatment significantly changed root-to-aerial or aerial-to-grain transfer of Cd compared to the BW-NT control (Figure 4C,D). There was a slight increase in aerial-to-grain transfer of Cd with green manure in the BW-NAS line, but this was not statistically different from the BW-NT control (*p*=0.06, Figure 4D). Transfer factors for Cu showed a significant increase in root-to-aerial transfer in BW-NAS with green manure compared to BW-NT with green manure (Figure S6-A). Transfer factors for Fe are not provided due to the presence of soil particles in the roots, despite careful washing, which led to Fe contamination.

**Figure 4.**
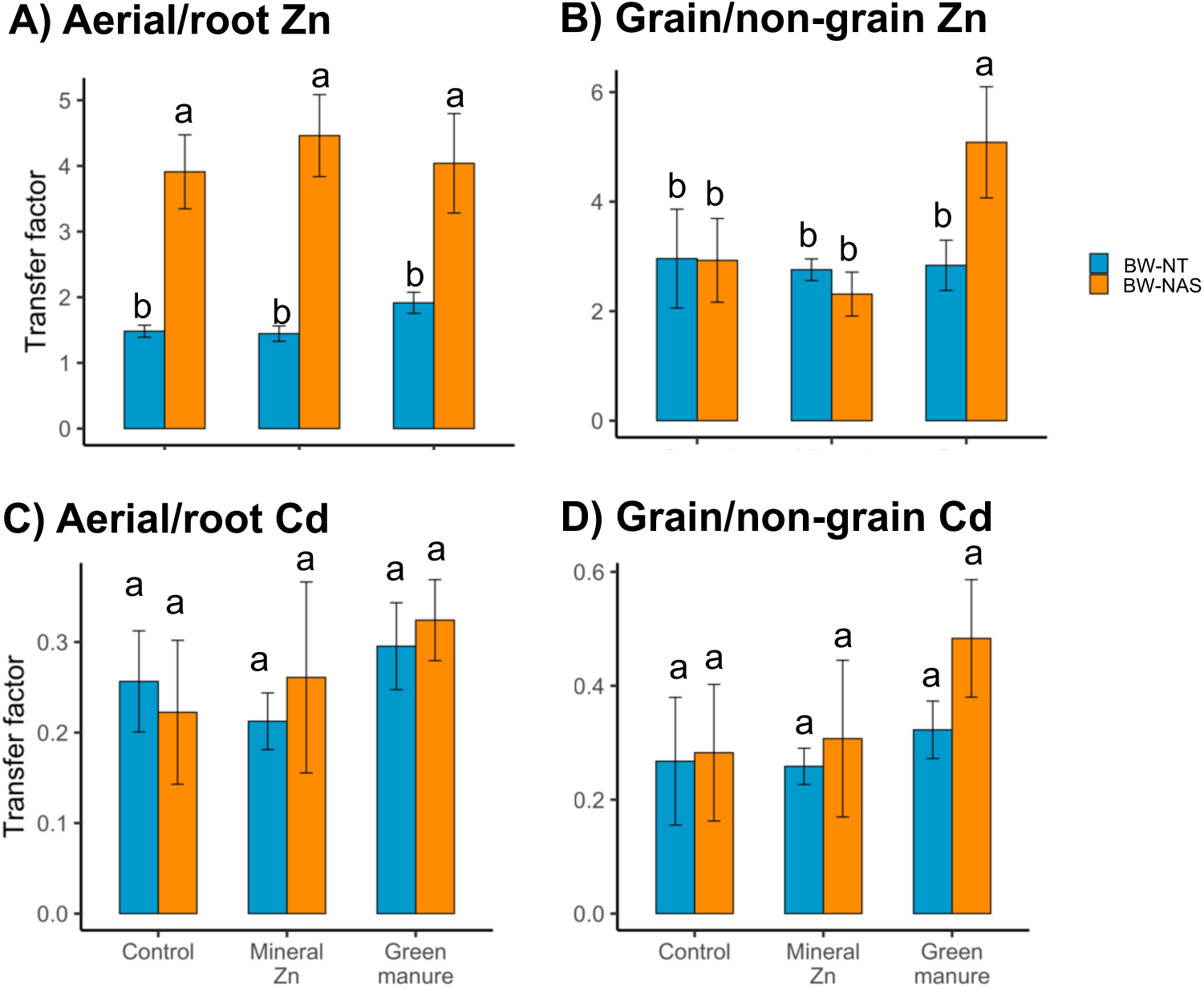
Transfer factors of Zn and Cd in non-transgenic (BW-NT) and *OsNAS2*-expressing (BW-NAS) wheat lines grown under three input treatments (control, mineral Zn, and sunflower green manure application). Values were determined from concentrations at maturity and include A) Zn ratio between aerial and root biomass, B) Zn ratio between grains and all other aerial biomass, C) Cd ratio between aerial and root biomass, and D) Cd ratio between grains and all other aerial biomass. Error bars indicate one standard deviation about the mean, of n=4 biological (pot) replicates. Lowercase letters indicate statistical differences calculated using generalized least squares modelling followed by emmeans posthoc tests. Significant *p*-values for the two-way ANOVA analyses based on generalized least squares output were as follows: A) line *p*<0.001; B) line *p*=0.03, inputs *p*<0.001, line*inputs *p*<0.001, and C) inputs *p*=0.04.

## Discussion

### Grain Zn increased due to combined effect of soil inputs and constitutive *OsNAS2* expression in wheat line

To our knowledge, this study is the first to investigate the combined effects of an *OsNAS2*-expressing wheat line and agronomic biofortification on Zn and Cd in wheat crops. To achieve this, we quantified grain concentrations, uptake, and internal translocation of Zn and Cd. Previous work has shown *OsNAS2* expression leads to increased Zn in cereal crops, mainly rice and barley (Masuda et al. 2009; Johnson et al. 2011; Lee et al. 2012; Kawakami and Bhullar 2020), but also in wheat (Singh et al. 2017; Beasley et al. 2022). We have confirmed the results of <u>Singh et al. (2017)</u> showing this for wheat (Figure 2A) and have demonstrated this in a non-artificial system (i.e. in real soil, as opposed to hydroponics or “artificial” model soil) with no discernable artefacts resulting from prior soil spiking (Note S3). As the *OsNAS2-*expressing line took up large amounts of Zn (∼25% of total uptake) between flowering and maturity (Table S7), it is likely that continuous uptake during grain filling contributed to higher Zn grain concentrations in BW-NAS than BW-NT, a mechanism proposed by Cakmak and Kutman (Kutman et al. 2012; Cakmak and Kutman 2018).

The increase in grain Zn and not Cd concentrations in the BW-NAS compared to BW-NT line can be explained by mechanisms of soil-to-plant and root-to-aerial transfer. Increased uptake of Zn but not Cd with in the *OsNAS2-*expressing line may have been due to increased production of the phytosiderophore DMA. Release of phytosiderophores via root exudation can solubilize previously unavailable soil species of Zn, allowing rapid uptake by plants (Clemens 2019). The importance of phytosiderophore-mediated Zn uptake in BW-NAS is further supported by a lack of concomitant increase in Cd uptake, as phytosiderophores have been shown to selectively increase uptake of metal micronutrients (Shenker et al. 2001; Meda et al. 2007; Puschenreiter et al. 2017; Banakar et al. 2017). We also observed a large increase in root-to-aerial transfer of Zn in *OsNAS2-*expressing wheat line as compared to the BW-NT in all input treatments (Figure 3A). Expression of *NAS* genes has been shown to increase root-to-aerial transfer of Zn in cereal crops, as both nicotianamine and DMA can increase symplastic mobility in roots and facilitate xylem loading (Suzuki et al. 2008; Deinlein et al. 2012; Haydon et al. 2012; Clemens et al. 2013). This increased transfer may have led to efficient removal of Zn from roots, shifting homeostasis and allowing further uptake from the soil. Yet, we did not observe increased root-to-aerial transfer of Cd, likely due to the higher affinity of Zn for low-molecular-weight chelating compounds containing N- and O-functional groups compared to Cd (Suzuki et al. 2007). Wang et al. (2013) reported that rice cultivars grown under high Fe conditions signal Zn deficiency by upregulating *OsZIP1* and *OsZIP4*, which mediate root Zn uptake and root-to-aerial translocation of Zn, respectively (Olsen and Palmgren 2014). This suggests that the increased Zn uptake observed in BW-NAS could also have been influenced by internal plant regulatory processes independent of *OsNAS2* expression, such as Zn-deficiency signaling triggered by enhanced Fe mobilization through phytosiderophore release. Notably our results differ from those of Hollmann et al. (2025), who reported that increased nicotianamine synthesis elevated the Cd accumulation in *Arabidopsis thaliana* grown in artificially contaminated soil containing 2 mg Cd kg⁻¹. Whether constitutive *OsNAS2* expression also increases grain Cd under substantially higher soil Cd concentrations, such as in contaminated systems, remains to be verified.

In this work, we demonstrated that combining an *OsNAS2*-expressing wheat line and input application led to greater increases in grain Zn than the additive effects of each strategy individually (Figure 2A). We also observed that the combination of BW-NAS and mineral Zn application led to a large increase in wheat uptake of Zn, i.e., two-fold higher than the expected additive increase (Figure 3A). Thus, our results strongly suggest that with combined BW-NAS and mineral Zn application, the synergistic increase in grain Zn was caused by increased uptake of Zn. Combined BW-NAS and green manure application led to increased transfer of Zn from non-grain aerial biomass to grains compared to BW-NT (Figure 4B), while wheat Zn uptake was equal to the expected additive effect (Figures 3A). Thus, increased transfer to grains explains the synergistic increase in grain Zn concentrations with combined BW-NAS and green manure compared to the BW-NT control.

We also observed increased grain Cd in the combined BW-NAS/green manure treatment compared to all other treatments (Figure 2B). One contributing factor may be a concentrating effect of reduced grain biomass compared to the BW-NT grown with green manure (Figure 1E). We also observed a non-significant trend towards increased aerial-to-grain transfer of Cd compared to the BW-NT control (*p=*0.06, Figure 4D), which may have further contributed to increased grain Cd. This is supported by the observation that the 17% reduction in grain biomass in BW-NAS/green manure treatment compared to BW-NT/green manure cannot fully explain the two-fold increase in grain Cd concentrations, nor the significantly higher concentrations compared to the BW-NT/control (whose grain biomass did not differ significantly from BW-NAS/green manure; Figure 1E). Thus, the concomitant increase could be because Cd can use the same membrane transporters as Zn (Bækgaard et al. 2010; Lin and Aarts 2012). Whether the observed increase in grain Cd is attributable to the single transgenic event used in the study cannot be sufficiently inferred from the current data and would require evaluation of additional independent *OsNAS2* lines in future work. Nevertheless, these observations highlight the importance of monitoring Cd when organic amendments are combined with genetic biofortification strategies.

### High fertilizer use efficiency of Zn and Cd with constitutive *OsNAS2* expression in wheat

The BW-NT line showed 5% Zn use efficiency with mineral Zn, which is comparable to previous works showing 2-10 % (Hippler et al. 2015; Mattiello et al. 2021; Yan et al. 2021). The use efficiencies of Zn and Cd from green manure were higher in this study compared to previous work. For Zn, we found 25% use efficiency from green manure in BW-NT (Figure 3C) compared to 1-16% in previous work (Aghili et al. 2014; Künzli et al. 2024). For Cd, we found 9 % use efficiency in BW-NT (Figure 3F), which is higher than the 2-7 % reported previously (Künzli et al. 2024). The higher green manure Zn and Cd use efficiencies in our study are likely due to the low pH of our soil, leading to increased Zn and Cd availability with organic amendment application compared to the alkaline conditions of these previous studies (Smolders and Mertens 2013; Mertens and Smolders 2013). The Zn use efficiencies in the BW-NAS line were very high compared to values reported previously (i.e., 15 % with mineral Zn, 40 % with green manure). This follows that BW-NAS likely has increased ability to solubilize and take up Zn from the soil compared to BW-NT, as discussed in the previous section. Our highly efficient wheat line (BW-NAS) was able to take up a much larger portion of Zn applied than previously reported, indicating the potential for much greater efficiency of soil Zn fertilization if combined with *OsNAS2-*expressing wheat line. Based on our above comparison with previous literature, the ability of BW-NAS to increase Zn use efficiency in alkaline, calcareous soil may be lower than in the moderately acidic soil used in this study; further investigation is required to confirm this.

### Concomitant increases in wheat Fe and Cu with combined genetic and agronomic biofortification

While our study was focused on Zn and Cd, we observed that the *OsNAS2-*expressing line and input application also affected Fe and Cu in wheat. The observed increase in grain Fe in BW-NAS compared to BW-NT (Figure 2C) follows previous studies showing that *NAS* expression leads to concomitant increases in cereal grain Zn and Fe (Singh et al. 2017; Kawakami and Bhullar 2018, 2020). This was likely via increased Fe uptake and transfer to grains facilitated by DMA and nicotianamine (Wirth et al. 2009; Wang et al. 2013; Ariga et al. 2014). When considered individually (i.e., without combination with inputs), BW-NAS did not show increased grain Cu compared to BW-NT (Figure 2D). Singh et al. (2017) found grain Cu concentration increased by a factor of 1.2 in the *OsNAS2* expressing wheat, which was low compared to their observed increases in grain Zn and Fe by a factor of 2. Thus, it seems the *OsNAS2* expressing line used in this study did not show a consistent, large increase in grain Cu as it does for Zn and Fe, which aligns with previous work (Haydon et al. 2012). Moreover, grain Cu concentrations measured in this study were much lower than values measured by <u>Singh et al. (2017)</u> (i.e. >5 mg Cu (kg grain)^−1^ in Singh et al. compared to <3 mg Cu (kg grain)^− 1^ in this study). This could indicate that constitutive *OsNAS2* expression only increases grain Cu in wheat when there is adequate availability of Cu. This is supported by the fact that we observed a significant increase in grain Cu with combined BW-NAS and green manure application. Green manure may have increased soil Cu via direct addition or by promoting solubilization of previously unavailable Cu species. This may explain the observed increase in water-soluble Cu (Figure S4-C), which is indicative of potentially higher availability. Furthermore, nicotianamine has been shown to be an important molecule for Cu transport (Takahashi et al. 2003). In the first discussion section, we postulate that combined BW-NAS and input application led to large increases in nicotianamine and DMA production. These increases may have led to the observed slight concomitant increases in grain Cu and the significant increase in Cu total uptake and root-to-aerial transfer in BW-NAS grown with green manure, compared to BW-NT in green manure (Figures S3-E, S6-A).

### Future work

This study provides a framework for future investigations of combined genetic biofortification with agronomic strategies such as mineral and organic input application. Our results indicate that nicotianamine and DMA likely play an important role in increasing grain Zn, Fe, and Cu while mitigating Cd in *OsNAS2* expressing wheat. Direct measurements are needed to confirm whether increased uptake and root-to-aerial transfer of Zn and Fe with *OsNAS2* expression resulted from increased nicotianamine and DMA production. This can be achieved via quantification of nicotianamine, DMA, and associated metals in the xylem sap, phloem sap, and root exudates of *OsNAS2* expressing wheat, using mass spectrometric techniques (Nishiyama et al. 2012; Oburger et al. 2014; Flis et al. 2016; Boiteau et al. 2018). Further investigation into the response of grain yield in *OsNAS2-*expressing wheat grown under organic amendments would be useful, as there was a slight decrease in grain mass per plant in BW-NAS compared to BW-NT in one of the treatment, i.e., the green manure. Since no difference in 1000 g weight was observed, future work should consider how changes in biomass allocation in *OsNAS2-*expressing wheat lines affects Zn and Cd concentrations in wheat grains. For the first time, we showed that combined *OsNAS2* expression and green manure application has the potential to increase grain Cd. This finding needs to be evaluated for a wide range of soil properties, especially in alkaline, calcareous soil. This is because green manure has been previously shown to facilitate solubilization of Cd in high pH soil (Grüter et al. 2018; Künzli et al. 2024), which could exacerbate the finding of the current study that combined *OsNAS2* expression and green manure increased wheat grain Cd compared to the BW-NT control. Finally, constitutive expression of *OsNAS2* has been shown to increase Zn bound to nicotianamine and DMA, which may also increase Zn availability in resulting food products (Lee et al. 2011). It needs to be investigated how our combined genetic and agronomic biofortification strategies affect allocation and chemical forms of Zn and Cd to edible grain biomass (Yan et al. 2025), and how this affects absorption of the nutrients when consumed.

## Conclusions

Our study demonstrates that genetic biofortification via an efficient, *OsNAS2-*expressing wheat line led to a greater increase in grain Zn compared to agronomic biofortification via input application to soil alone. Importantly, combining the *OsNAS2* expressing line and input application lead to synergistic increases in grain Zn that surpassed the expected additive effects of each strategy individually. These results highlight the strong potential of integrating genetic and agronomic approaches to achieve nutritionally meaningful improvements in grain micronutrient content. Concomitant increases in grain Fe and Cu were also observed, especially when combining the *OsNAS2-*expressing wheat line and green manure application. However, the combined *OsNAS2*/green-manure treatment also resulted in elevated grain Cd relative to the non-transgenic control, underscoring the need for careful monitoring of grain Cd concentrations when organic amendments are paired with genetic biofortification strategies. Further work is required to determine whether such Cd increases occur across a broader range of soils associated with low Zn availability, including high-pH and saline environments. Overall, our findings support the development of integrated biofortification strategies that combine genetic enhancement with targeted agronomic practices to maximize Zn and Fe concentrations in wheat grains while ensuring safe levels of Cd.

## Statements and declarations

### Author contributions

**JB**: conceptualization, validation, formal analysis, investigation, writing – original draft, visualization, project administration. **JT**: conceptualization, methodology, investigation, writing – reviewing & editing, supervision, funding acquisition. **LHEW**: conceptualization, writing – reviewing & editing, supervision, funding acquisition. **NKB:** conceptualization, writing – reviewing & editing. **CM**: investigation, writing – reviewing & editing. **SAY**: investigation, writing – reviewing & editing. **MW**: conceptualization, investigation, writing – reviewing & editing (primary input), supervision, funding acquisition. **EF**: conceptualization, writing – reviewing & editing (primary input), supervision, project administration, funding acquisition.

## Acknowledgements

We are grateful to Monika Macsai, who ensured healthy plant growth throughout the experiment. We wish to thank Monika, Federica Tamburini, and Astrid Oberson who contributed to isotope-labeled green manure production even when the university was under covid lockdown. We are grateful to Camille Rubeaud, whose preliminary experiment with the BW-NT and BW-NAS lines helped inform our experimental design. Thanks are due to Monika, Federica, and Laurie Mauclaire-Schoenholzer for their support in experimental setup, harvesting, and analyses. Special thanks go to Laurie for her help with double-distilling the acids used in this experiment and her analyses of dissolved organic carbon. We also thank Federica for her kind help with CNS analyses. We are grateful to Lisa Konrad, Fernando Brito, Florent Leyvraz, and Robin Krischer for support with analyzing samples and washing roots. We thank Geremia Pellegri for his help maintaining the wheat crops and monitoring their growth during the experiment. Many thanks to Natalie Gsteiger at Eawag, and all staff and apprentices from AuA, for helping us characterize nitrate in soil water extracts. We are also very grateful to Corey Archer for his aid in constructing the columns used for isotope purifications and his helpful advice on decreasing zinc contamination in our samples. RT-PCR data produced and analyzed in this paper were generated in collaboration with the Genetic Diversity Centre of ETH Zurich. Finally, our deepest gratitude goes to the Group of Plant Nutrition and the Group of Inorganic Environmental Geochemistry at ETH and Eawag for all their help and support during this project.

