## Supplementary Information for "Combining agronomic biofortification with a Zn-efficient wheat line synergistically increases grain Zn concentrations with varied effects on Cd in wheat grains"

**Affiliations:**

^#a^Institute of Earth Surface Dynamics, University of Lausanne, Lausanne, Switzerland

^#b^Federal Institute of Metrology METAS, Bern-Wabern, Switzerland

^#c^Intagris AG, Regensdorf, Switzerland

^#d^Puregene AG, Zeiningen, Aargau, Switzerland

**Table S1. Properties of soil (0-20 cm, topsoil) and sunflower green manure.** Error indicates one standard deviation calculated from three processed replicates.

| **Property** | **Unit** | **Soil** | **Green manure** |
| --- | --- | --- | --- |
| pH^a^ |  | 5.5* | 6.7 ± 0.0 |
| C^b^ | g (kg)^-1^ | 16 ± 1 | 346 ± 8 |
| N^b^ | g (kg)^-1^ | 1.7 ± 0.1 | 56 ± 2 |
| N-NH_4_ ^c^ | mg (kg)^-1^ |  | 91 |
| N-NO_3_ ^d^ | g (kg)^-1^ |  | 1.1 |
| S^b^ | mg (kg)^-1^ | 0.208 ± 0.001 |  |
| DTPA-Zn^e^ | mg (kg)^-1^ | 0.8 ± 0.2 |  |
| DTPA-Cd^e^ | mg (kg)^-1^ | 0.09 ± 0.03 |  |
| Total Zn^f^ | mg (kg)^-1^ | 95.2 ± 0.1 | 11 |
| Total Cd^f^ | mg (kg)^-1^ | 0.89 ± 0.01 | 0.89 |
| Clay^g^ | g (kg)^-1^ | 192* |  |
| Silt^g^ | g (kg)^-1^ | 633* |  |
| Sand^g^ | g (kg)^-1^ | 175* |  |
| WHC^h^ | g (kg)^-1^ | 320 ± 7 |  |
| CEC^i^ | meq (100 g)^-1^ | 13.5* |  |
| ^a^Measured in water, 1:2.5 for soil (Agroscope 2020a) and 1:50 for green manure | | | |
| ^b^Elemental analyzer, Vario PYRO cube, Elementar GmbH, Germany | | | |
| ^c^Berhelot’s reaction with Agilent Cary 60 Spectrophotometer (Searle 1984) | | | |
| ^d^Ion chromatography via metrohm 930 Compact IC Flex with chemical suppression | | | |
| ^e^Diethylenetriaminepentaacetic acid extraction (Lindsay and Norvell 1978) | | | |
| ^f^Acid digestion (HF for soil, HNO_3_ for green manure) | | | |
| ^g^Gravimetric analysis (Agroscope 2020b) | | |  |
| ^h^Water-holding capacity (Agroscope 2020c) | | |  |
| ^i^Cation exchange capacity (Agroscope 2020d) | | |  |
| *Measurement performed by Sol Conseil, Gland, Switzerland | | | |

**Table S2. Modifications to full-strength Hoagland nutrient solution used in production of ^70^Zn and ^106^Cd-labeled green manure.** Green manure was produced by growing sunflower (*Helianthus annuus*) hydroponically. Nutrient solutions were prepared in MilliQ water following the recipe for full-strength Hoagland solution two (Hoagland and Arnon 1950). The Hoagland solution was modified in the quantity and form of Zn, and Cd was also added, as shown in the table below. Values for Zn and Cd added at natural abundance (NA) and as ^70^Zn and ^106^Cd spikes are provided on separate lines in the table below.

| **Element** | **Element concentration, mg (L)^-1^** | **Salt** |
| --- | --- | --- |
| Zn (NA) | 0.05 | Zn(NO_3_)_2_ |
| ^70^Zn | 0.05 | ^70^Zn(NO_3_)_2_ |
| Cd (NA) | 0.01 | Cd(NO_3_)_2_ |
| ^111^Cd | 0.01 | ^111^Cd(NO_3_)_2_ |

**Table S3. Nutrient solution for production of new generation of wheat seeds (Bobwhite cultivar, non-transgenic and *OsNAS2-*expressing lines).** Wheat was fertilized approximately every two weeks between sowing and grain filling, for a total of eight fertilization events. Each fertilization included the listed elements at the provided application rate.

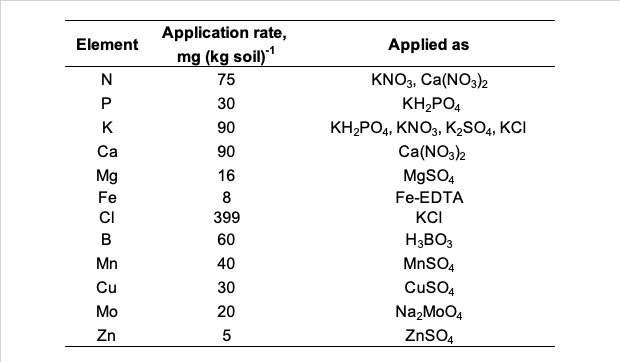

**Note S1**. **Further explanation of Zn and Cd addition to soil prior to pot experiment.**

In parallel to the experiment presented in this manuscript (pot experiment 2; hereafter “PE 2”), we performed a wheat-growth pot experiment to determine the effect of organic amendment chemical composition on Zn and Cd uptake in wheat (pot experiment 1; hereafter “PE1”) (Bachelder et al. 2026). PE 1 used the non-transgenic Bobwhite wheat line, i.e. “BW-NT” in this paper. The details of PE 1 can be found in our previous manuscript (Bachelder et al. 2026). PE 1 involved application of manure and compost, which cannot be easily spiked with isotopes in a homogeneous way (Fuchs et al. 2023). Thus, we performed the indirect spiking method, in which spikes of ^70^Zn and ^106^Cd were added to the soil, allowed to equilibrate with the available soil pool for twelve weeks, and used to distinguish the soil- and amendment-derived Zn and Cd, as explained in detail in Bachelder et al. (2026). To ensure comparability between the results of PE 1 and PE 2, we performed the same spiking procedure in PE 2 using Zn and Cd at natural abundance. For isotope tracing in PE 2, we applied ^70^Zn- and ^106^Cd-spiked inputs to perform the higher-precision direct labeling method (Dürr-Auster et al. 2019). We also included two treatments to further evaluate the comparability between the experiments. First, we included a treatment of native soil only, in which we grew wheat in soil that did not go through the Zn and Cd spiking and twelve-week equilibration process. In PE 1, we also included a treatment of unlabeled sunflower with the following composition: pH = 6.3 ± 0.1, carbon = 338 ± 14 g (kg) ^-1^, nitrogen = 338 ± 14 g (kg) ^-1^, Zn = 12 ± 1 mg (kg)^-1^, Cd = 1.0 ± 0.1 mg (kg)^-1^. The unlabeled green manure was produced in parallel to the ^70^Zn- and ^106^Cd-spiked sunflower applied in this experiment. Hence, we could compare between PE 1 and PE 2 for the effects of the control, mineral Zn, and green manure treatments on Zn and Cd in the non-transgenic Bobwhite line using both indirect (PE 1) and direct (PE 2) isotope labeling.

**Table S4. Basal nutrient solution composition and input application rates for wheat-growth pot experiment.** Non-transgenic (BW-NT) and *OsNAS2*-expressing (BW-NAS) wheat lines were grown with three treatments (control, mineral Zn, and sunflower green manure application). The nutrient solution and treatments were mixed into the soil three days prior to sowing. Calcium nitrate was adjusted in the green manure treatment to maintain a constant application rate of mineral N.

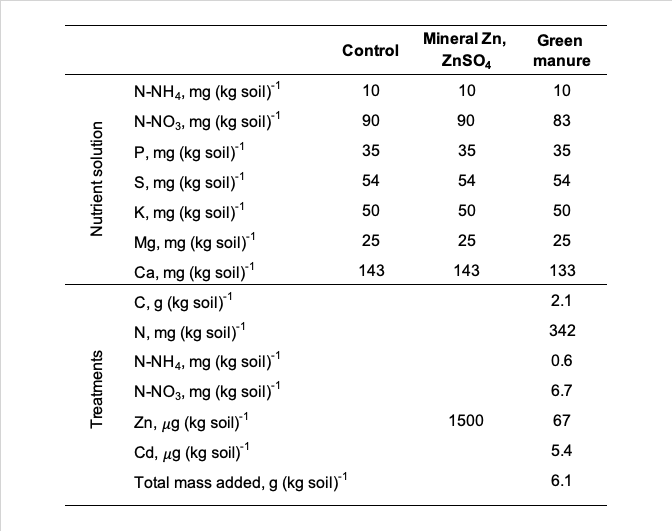

**Table S5. Quality control for ICP-MS/MS quantification of solid-phase elemental composition.** Certified values are from WEPAL 2020 measurement of Raye Grass (*Lolium perene*). The collision gases and isotopes analyzed are provided, with “-->” indicating the isotope of interest first followed by the isotope measured. Recovery is also provided, i.e. the percentage of the certified concentration recorded by our analyses. Results are for n=8 replicates.

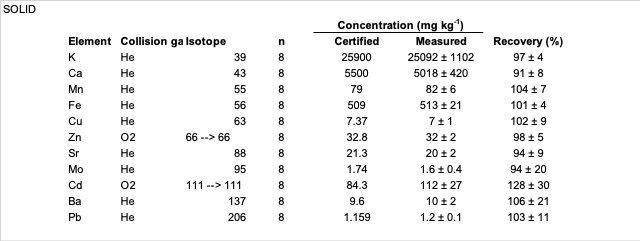

**Table S6. Quality control for ICP-MS/MS quantification of aqueous samples.** Analysis was performed using standard reference material 1643f (“trace elements in water”) from the national institute of standards and technology. The collision gases and isotopes analyzed are provided, with “-->” indicating the isotope of interest followed by the isotope measured. Recovery is also provided, i.e. the percentage of the certified concentration recorded by our analyses. Results are for n=6 replicates.

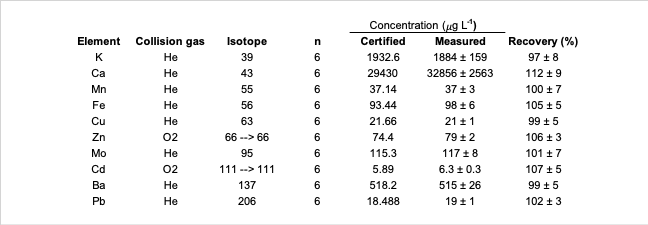

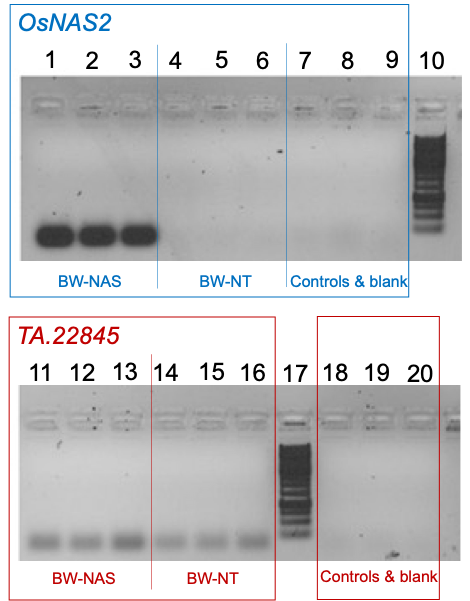

**Figure S1. Gel electrophoresis image of the amplification of the genes *OsNAS2* and *Ta.22845* in transgenic (BW-NAS) and wildtype (BW-NT) Bobwhite wheat lines.** Wheat leaves were sampled at flowering and immediately flash-frozen in liquid nitrogen. RNA was extracted and converted into cDNA prior to the amplification of *OsNAS2* gene using polymerase chain reaction (PCR). Samples 1-3 show the *OsNAS2* gene in the BW-NAS line. Samples 4-6 show the *OsNAS2* gene in the BW-NT line. Samples 7 and 8 are negative controls of the *OsNAS2* gene, i.e. PCR performed with cDNA from BW-NAS and BW-NT lines, respectively, obtained using the same reverse transcription protocol but without reverse transcriptase (therefore they should not contain any cDNA). Sample 9 is a water blank of the *OsNAS2* gene containing only PCR master mix and no cDNA. Samples 11-13 show the *Ta.22845* reference gene in the BW-NAS line. Samples 14-16 show the *Ta.22845* reference gene in the BW-NT line. Samples 18 and 19 are negative controls of the *Ta.22845* reference gene, i.e. PCR performed with cDNA from BW-NAS and BW-NT lines, respectively, obtained using the same reverse transcription protocol but without reverse transcriptase. Sample 20 is a water blank of the *Ta.22845* reference gene containing only PCR master mix and no cDNA. Samples 10 and 17 are the DNA ladder (GeneRuler 1 kb Plus DNA Ladder, ThermoFisher).

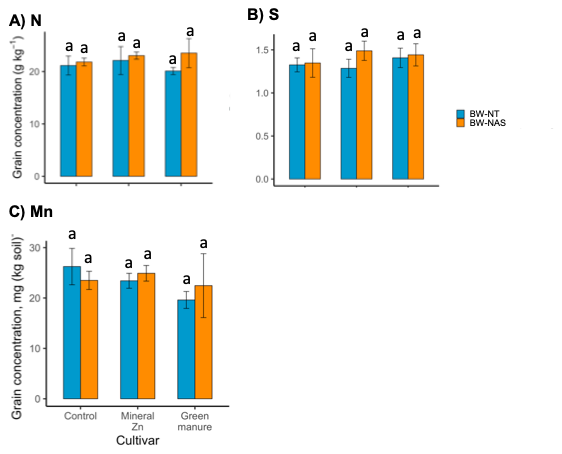

**Figure S2. Elemental grain concentrations in *OsNAS2* expressing Bobwhite (BW-NAS) and non-transgenic (BW-NT) wheat lines grown under three input treatments.** Panels included A) nitrogen, B) sulfur, and C) manganese measured in non-transgenic and *OsNAS2*-expressing wheat lines grown under three input treatments (control, mineral Zn, and green manure application). Error bars indicate one standard deviation about the mean, of n=4 biological (pot) replicates. Lowercase letters indicate statistical differences calculated using generalized least squares modelling followed by emmeans posthoc tests. Significant *p*-values for the two-way ANOVA analyses based on generalized least squares output were as follows: A) line *p*= 0.02; C) inputs *p*=0.03.

**Table S7. Total uptake of Zn, Fe, and Cd at flowering (H1) and maturity (H2).** Data presented include *OsNAS2* expressing Bobwhite (BW-NAS) and non-transgenic (BW-NT) wheat lines grown under three input treatments: control, mineral Zn, and green manure application. Values were determined as the total elemental contents in roots and all aerial biomass (stem + leaves at flowering, stem + leaves + grains at maturity). The average (av) and standard deviation (sd) of n=4 biological (pot) replicates is provided. Lowercase letters indicate statistical differences calculated using a paired t-test comparing H1 and H2 for each wheat line/input treatment (i.e., only values within black boxes are compared). This allowed evaluation of whether there was a statistical difference (i.e., *p*-value < 0.05) between the total uptake at H1 compared to H2, for each treatment. Instances where there was a statistical difference are highlighted with bold text and yellow shading.

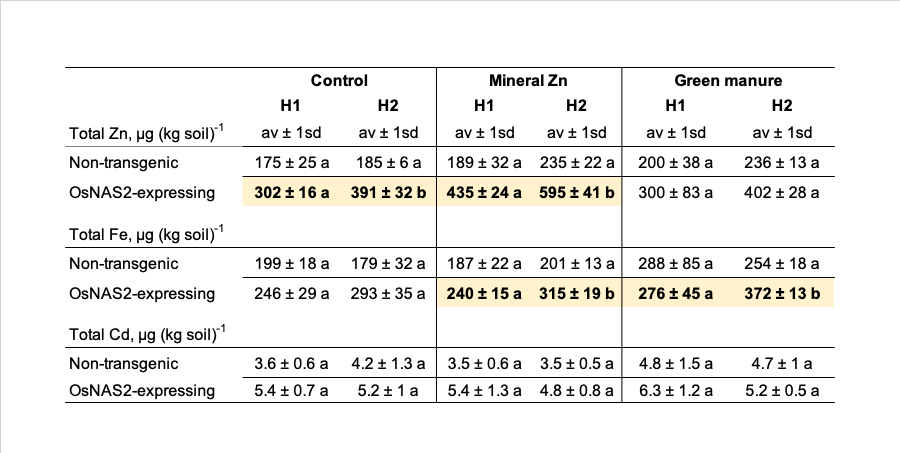

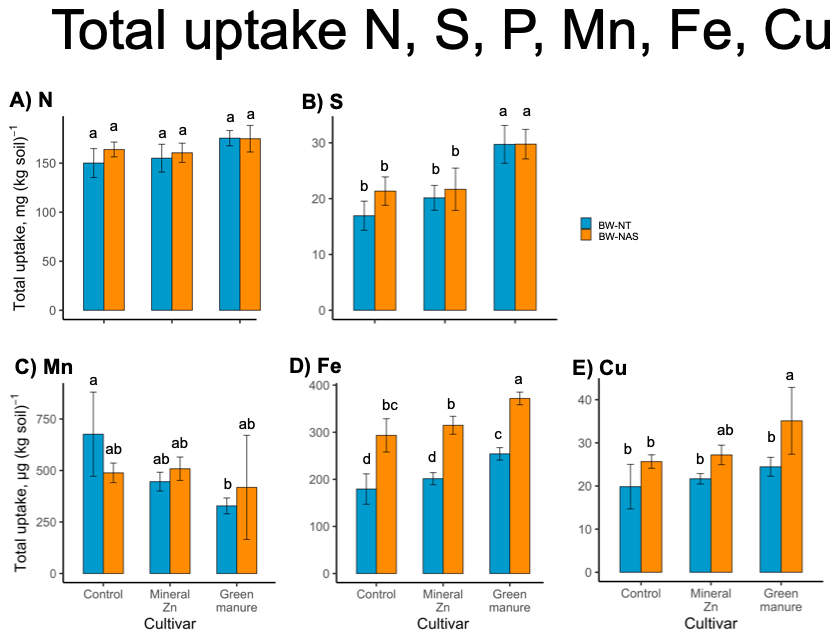

**Figure S3. Total uptake of A) nitrogen, B) sulfur, C) manganese, D) iron, and E) copper in wheat at maturity.** Data presented include *OsNAS2* expressing Bobwhite (BW-NAS) and non-transgenic (BW-NT) wheat lines grown under three input treatments: control, mineral Zn, and green manure application. Values were determined for roots and all aboveground biomass (stem + leaves and grains). Error bars indicate one standard deviation about the mean, of n=4 biological (pot) replicates. Lowercase letters indicate statistical differences calculated using generalized least squares modelling followed by emmeans posthoc tests. Significant *p*-values for the two-way ANOVA analyses based on generalized least squares output were as follows: A) inputs *p*=0.002; B) inputs *p*<0.001; C) inputs *p*=0.02; D) line *p*<0.001, inputs *p*<0.001; E) line *p*<0.001, inputs *p*=0.001.

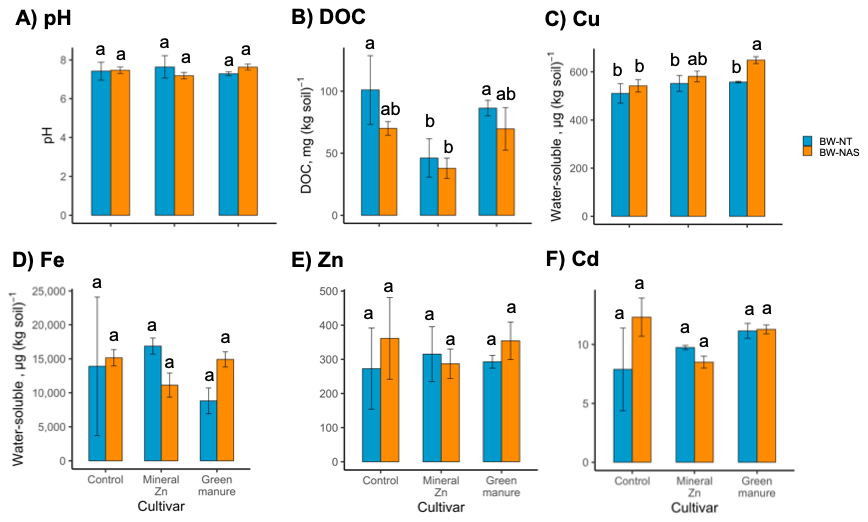

**Figure S4. Soil properties measured at flowering for *OsNAS2* expressing Bobwhite (BW-NAS) and non-transgenic (BW-NT) wheat lines were grown under three input treatments.** Input treatments included control, mineral Zn, and green manure application. Data includes A) pH, B) dissolved organic carbon (DOC), and water-soluble pools of C) Cu, D) Fe, E) Zn, and F) Cd. Values were determined at maturity for roots and all aboveground biomass (stem, leaves, and grains). Error bars indicate one standard deviation about the mean, of n=4 biological (pot) replicates, except for water-soluble pools (D-F) where only three replicates were included. Lowercase letters indicate statistical differences calculated using generalized least squares modelling followed by emmeans posthoc tests. Significant *p*-values for the two-way ANOVA analyses based on generalized least squares output were as follows: A) line*inputs *p*=0.04, B) line *p*=0.002, inputs *p*<0.001; C) line *p*<0.001, inputs *p*<0.001; F) line*inputs *p*=0.01.

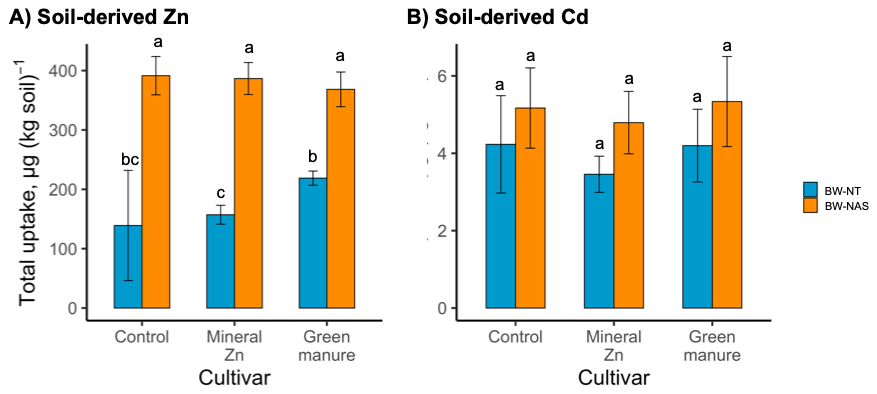

**Figure S5. Soil-derived A) Zn and B) Cd measured in wheat biomass at maturity.** Values are from *OsNAS2* expressing Bobwhite (BW-NAS) and non-transgenic (BW-NT) wheat lines grown under three input treatments (control, mineral Zn, and green manure application). Values were determined for roots and all aboveground biomass (stem, leaves and grains). Error bars indicate one standard deviation about the mean, of n=4 biological (pot) replicates. Lowercase letters indicate statistical differences calculated using generalized least squares modelling followed by emmeans posthoc tests. Significant *p*-values for the two-way ANOVA analyses based on generalized least squares output were as follows: A) line *p*<0.001, line*inputs *p*=0.003; B) line *p*=0.004.

**Table S8**. **Grain concentrations and Zn and Cd use efficiencies between pot experiment 1 (PE 1; Bachelder et al. 2026) and this experiment (pot experiment 2, i.e. PE 2).** In PE 1, wheat was grown on soil spiked with 200 μg ^70^Zn (kg soil)^-1^ and 2 μg ^106^Cd (kg soil)^-1^. In PE 2, wheat was grown on soil spiked with 200 μg Zn (kg soil)^-1^ and 2 μg Cd (kg soil)^-1^, i.e. Zn and Cd at natural abundance. The data presented are for non-transgenic Bobwhite wheat grown without Zn fertilization (no Zn, or “control” in main text), with mineral Zn fertilization (1.5 mg Zn applied per kg soil), and with green manure (6 g sunflower applied per kg soil). The green manure treatment from PE 1 was not published due to high variability of the isotope tracing results. To determine the effect of soil spiking on grain Zn and Cd, a separate treatment of “native soil only” was also included containing no Zn and Cd spike and now twelve-week incubation prior to the pot experiment. Use efficiency was determined as the percentage of Zn and Cd applied at the beginning of the experiment that was quantified in plant biomass at maturity, using isotope tracing. Error is provided as one standard deviation about the mean of n=4 biological (pot) replicates.

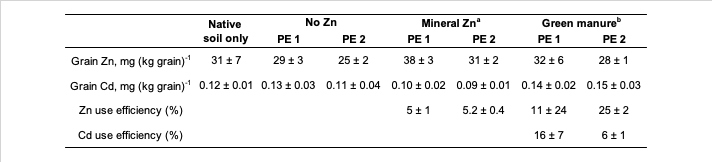

^a^Mineral Zn was applied as ZnSO_4_ containing Zn at natural abundance in PE 1 and as 10% ^70^Zn(NO_3_)_2_, 90% ZnSO_4_ in PE 2.

^b^Green manure was applied as sunflower, which contained Zn and Cd at natural abundance in PE 1 and was ^70^Zn- and ^106^Cd-spiked in PE 2.

**Note S2. Comparability between pot experiment 1 (PE 1; Bachelder et al. 2026) and this experiment (pot experiment 2, i.e. PE 2).**

The two experiments are described in detail in the caption of Table S8. As presented in Table S8, both PE 1 and PE 2 showed grain Zn and Cd concentrations that were similar to wheat grown in native soil. PE 2 grain Zn concentrations tended to be lower than in PE 1, but showed similar trends (i.e., mineral Zn > green manure> no Zn). Overall, adding Zn and Cd spikes in PE 1 and PE 2 produced no observable change in grain Zn and Cd compared to the native soil. The Zn use efficiency for the mineral Zn treatment was very similar in PE 1 and PE 2, with lower standard deviation in PE 2 (5.2 ± 0.4) compared to PE 1 (5 ± 1). The higher precision in PE 2 compared to PE 1 was due to the use of the direct spiking method, as outlined by previous work (Dürr-Auster et al. 2019). With green manure, the variability in use efficiency calculations was also much lower in PE 2 compared to PE 1, with 25 ± 2 % compared to 11 ± 24 % for Zn and 6 ± 1 % compared to 16 ± 7 % for Cd, respectively. This was also due to the use of direct spiking in PE 2, versus indirect spiking in PE 1. For both mineral Zn and green manure, the values of PE 2 use efficiencies fall in the range of those observed in PE 1.

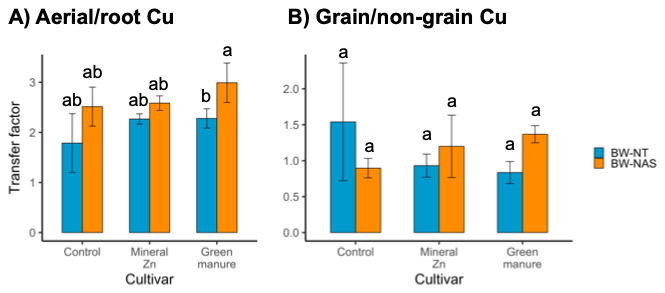

**Figure S6. Transfer factors calculated from concentrations of Cu in wheat biomass at maturity.** Values are from *OsNAS2* expressing Bobwhite (BW-NAS) and non-transgenic (BW-NT) wheat lines grown under three input treatments (control, mineral Zn, and green manure application). Panels include A) Cu ratio between aboveground and belowground biomass and B) Cu ratio between grains and all other aboveground biomass. Lowercase letters indicate statistical differences calculated using generalized least squares modelling followed by emmeans posthoc tests. Significant *p*-values for the two-way ANOVA analyses based on generalized least squares output were as follows: A) line *p*<0.001, inputs *p*=0.03; B) line *p*=0.03, inputs *p*<0.001, line*inputs *p*<0.001.

**Note S3. Evaluation of experimental artefacts resulting from addition of natural-abundance Zn and Cd to soil prior to pot experiment.**

We added 200 μg Zn and 2 μg Cd to the soil twelve weeks prior to the start of our experiment. This is a large amount of element compared to our available pools, as it is equal to ~100 % of Zn and ~50 % of Cd taken up by plants (see main text Figure 2A,C). Thus, we must consider whether our prior soil spike led to a significant increase in wheat uptake of Zn and Cd. In parallel to the experiment presented in this paper (PE 2), we performed a wheat-growth pot experiment to determine the effect of organic amendment chemical composition on Zn and Cd uptake in wheat (PE 1) (Bachelder et al. 2026). In PE 1, we added 200 μg ^70^Zn and 2 μg ^106^Cd in exactly the same manner as PE 2, allowing us to trace (and quantify) the spike in the plant biomass. In PE 1, the abundance of ^70^Zn in the control treatment was 7%, compared to 0.6% at natural abundance. Hence, if the plant took up 180 μg Zn (kg soil)^-1^, ~11 μg Zn (kg soil)^-1^ of this was derived from the added Zn, and in wheat grains containing 30 mg Zn (kg grain)^-1^, this would include ~1.9 mg derived from the added Zn. In PE 1, the abundance of ^106^Cd in the control was 2.5 %, compared to 1.25 % at natural abundance. Hence, if the plant took up 4 μg Cd (kg soil)^-1^, ~0.05 μg Cd was derived from the Cd spike, and in wheat grains containing 100 μg Cd (kg grain)^-1^, this corresponds to ~1.3 μg Cd (kg grain)^-1^ derived from the Cd spike. The quantities derived from the added Zn and Cd are similar to one standard deviation about the mean in our data (Table S8, main text Figure 2-A,C), meaning they are within the range of variability between our experimental (pot) replicates. Thus, we conclude the spike did not greatly affect Zn and Cd uptake and translocation to grains in this experiment.

Our conclusion that the spike did not greatly affect Zn and Cd uptake in wheat is supported by the fact that the grain concentrations of Zn and Cd were similar to the concentrations in wheat grown in soil with no spike addition (Table S8). The high variability in grain Zn in the test with no spike addition was likely due to the lack of mixing and equilibration of this treatment for twelve weeks prior to the experiment, as was performed for all treatments in PE 1 and PE 2. This supports our choice to process all experimental (pot) replicates in the same manner during our experiments, to decrease variability and increase the comparability of the experiments.
